# Multiple-gene targeting and mismatch tolerance can confound analysis of genome-wide pooled CRISPR screens

**DOI:** 10.1101/387258

**Authors:** Jean-Philippe Fortin, Karen E. Gascoigne, Peter M. Haverty, William F. Forrest, Michael R. Costa, Scott E. Martin

## Abstract

Genome-wide loss-of-function screens using the CRISPR/Cas9 system allow the efficient discovery of cancer cell vulnerabilities. While several studies have focused on correcting for DNA cleavage toxicity biases associated with copy number alterations, the effects of sgRNAs co-targeting multiple genomic loci in CRISPR screens have not been discussed yet. In this work, we analyze CRISPR essentiality screen data from 391 cancer cell lines to characterize biases induced by multi-target sgRNAs. We investigate two types of multi-targets: on-targets predicted through perfect sequence complementarity, and off-targets predicted through sequence complementarity with up to two nucleotide mismatches. We found that the number of on-targets and off-targets both increase sgRNA activity in a cell line-specific manner, and that existing additive models of gene knockout effects fail at capturing genetic interactions that may occur between co-targeted genes. We use synthetic lethality between paralog genes to show that genetic interactions can introduce biases in essentiality scores estimated from multi-target sgRNAs. We further show that single-mismatch tolerant sgRNAs can confound the analysis of gene essentiality and lead to incorrect co-essentiality functional networks.

## 1 Introduction

A central goal of functional genomics is to understand the complex relationship between the genotype and the phenotype of a given organism. Genome-scale forward genetic screens are powerful and unbiased experiments that help identify gene function and relationships between gene disruption and disease. In such screens, gene perturbations such as mutation or expression dysregulation are first introduced in cells or organisms, then phenotypes of interest in the cell or organism population are identified, and finally selected phenotypes are linked back to gene perturbations to establish causality. Genome-wide pooled screens exploiting the RNA interference (RNAi) pathway have significantly helped with identifying and prioritizing therapeutic targets in cancer and other diseases; several loss-of-function genome-wide RNAi screens across dozens of cancer cell lines have been conducted to study cancer genetic vulnerabilities as well as identify genes that are essential across cell lines [Tsherniak et al., 2017, Cowley et al., 2014, McDonald III et al., 2017, Nijhawan et al., 2012]. However, the analysis and interpretation of results obtained from RNAi screens can be confounded by ubiquitous off-target effects and incomplete knockdown.

The discovery of the CRISPR/Cas9 genome editing system and its application to functional screens have revolutionized the field by minimizing many of the challenges and limitations observed in RNAi screens [Shalem et al., 2015]. Following their previous effort in identifying cancer vulnerabilities using RNAi pooled screens [Tsherniak et al., 2017], the Broad Institute has been performing genome-wide pooled CRISPR knock-out screens across several hundreds of genomically characterized cancer cell lines [Meyers et al., 2017] using the Avana guide library [Doench et al., 2016]. The screening effort using both RNAi and CRISPR technologies is referred to as Project Achilles. To date, CRISPR screening data from 391 cell lines have been released and are available to download from the Project Achilles portal (https://portals.broadinstitute.org/achilles).

Early on, it was observed that variation in genomic copy number (CN) was impacting growth measurements in loss-of-function CRISPR screens in a gene-independent manner [Wang et al., 2015, Aguirre et al., 2016, Munoz et al., 2016]. Specifically, guides targeting amplified regions create a large number of DNA double-strand breaks (DSBs) resulting in a loss of cell viability, often referred to as “cleavage toxicity”. If not taken into account, gene-independent CN effects can lead to an increased number of false positives. In Meyers et al. [2017], the authors propose to delineate gene-knockout effects from CN effects in the Achilles dataset by concurrently modeling both effects in a linear regression framework; the algorithm is called CERES. In particular, the CN effects are captured in a cell-specific manner using linear splines, allowing for cell line-specific amplifications and deletions. A CN-adjusted essentiality score (CERES score) is estimated for each single-guide RNA (sgRNA) and for each cell line. Among others, CERES scores can be further analyzed to discover cancer vulnerabilities and gene essentiality.

Designing sgRNAs that uniquely map to the genome can be challenging, especially for genes sharing high homology with other genomic loci, either in coding or non-coding regions. In the Avana library, a number of sgRNAs are annotated to target multiple genes through perfect sequence complementarity between the sgRNA’s protospacer sequence and genomic DNA – we refer to such guides as “multi-target” guides. The CERES model attempts to account for multi-target effects by modeling the log-fold change (LFC) of a multi-target guide as a linear combination of knockout effects from the set of genes targeted by the guide; the model assumes that gene knockout effects are additive. For instance, for a guide targeting two genes, this assumes that the phenotypic effects of a double knockout is the sum of the individual gene knockout phenotypic effects. As a limitation, genetic interactions such as synergy, synthetic lethality, genetic buffering, and epistasis, cannot be appropriately captured by such a model. Additionally, while the CERES model attempts to account for multi-target effects, off-target effects caused by mismatch tolerance between the sgRNA’s protospacer sequence and genomic DNA have not been considered in the Achilles dataset.

In this work, we investigate the impact of multi-target effects on estimating cancer cell dependencies, as well as the impact of off-target effects caused by mismatch tolerance in sgRNA-DNA binding. We take advantage of the Achilles CRISPR screening data, by far the most comprehensive CRISPR screening effort to date, to generalize our findings across cell lines and sgRNAs. First, we show that the number of on-targets of a particular sgRNA can dramatically increase the sgRNA essentiality score in a non-additive fashion. To illustrate this, we consider guides in the Avana library that co-target paralogs *MYL12A* and *MYL12B*, and show that an additive model cannot capture the synthetic lethal interaction observed in a subset of cell lines in which the redundant third paralog *MYL9* is not expressed. We also show that off-target effects caused by single-mismatch sgRNA-DNA alignments can cause spurious associations between cell lineage and gene knockout. As an example, we found that several cell lines are unexpectedly reported as being dependent on *SOX9* despite the observation that *SOX9* is not expressed in these cell lines. We present evidence that off-target effects caused by single-mismatch tolerance are likely responsible for these inconsistent results. We provide gene-level summaries of on-target and off-target alignments in the Avana library to help flag and interpret genes with unexpected essentiality scores.

## 2 Material and Methods

### 2.1 Datasets

#### Achilles CRISPR

From the Achilles data portal (https://portals.broadinstitute.org/achilles), we downloaded CERES scores for 391 cell lines across 17,655 genes (file: `gene_effect.csv`). We also downloaded guide-level log-fold changes (`logfold_change.csv`); processing of these data is further discussed below.

#### Achilles RNAi

From the Achilles data portal (https://portals.broadinstitute.org/achilles), we downloaded gene-level DEMETER scores for 501 cell lines across 17,098 genes (file: `ExpandedGeneZSolsCleaned.csv`).

#### CCLE datasets

From the Cancer Cell Line Encyclopedia (CCLE) portal (https://portals.broadinstitute.org/ccle), we downloaded rpkm-level RNA-Seq data (file: `CCLE_RNAseq_081117.rpkm.gct`) and gene-level relative copy number data (file: `CCLE_copynumber_byGene_2013-12-03.txt`). We applied the transformation log_2_(rpkm + 1) to the RNA-Seq data.

### 2.2 sgRNA sequence alignments

For alignment, we considered the 72,787 guides in the Avana library for which LFCs were available in the portal. We used bowtie (v.1.2.2, [Langmead et al., 2009]) to align guide sequences to the Human genome assembly GRCh38, allowing up to 2 mismatches between the reference sequence (DNA) and the sgRNA’s protospacer sequence (bowtie with options -v 2 -k 10000). Using the R package BSgenome [Pagès, 2016] we filtered out alignments that did not have the canonical NGG PAM site, for a final total of 235,133 valid alignments. Using the comprehensive gene annotation from GENCODE v28 [Harrow et al., 2012], we added a gene and exon annotation for all alignments. Sequence alignments are provided in Supplementary File 1.

### 2.3 Processing of guide-level log-fold changes

We obtained guide-level raw LFCs from the Achilles portal (`logfold_change.csv`). As detailed in Meyers et al. [2017], fold changes were first calculated by dividing sample read counts by their representation in the starting plasmid DNA library (pDNA). As in Meyers et al. [2017], we normalized LFCs for each cell line replicate by centering the distribution using the median LFC value, and then dividing by the median absolute deviation (MAD). Since we are interested in visualizing and analyzing LFCs, we further scaled LFCs by the absolute average LFC value across cell lines for guides targeting essential genes (212 genes total, [Hart et al., 2014]), such that a value of −1 roughly indicates essentiality. We then averaged normalized LFCs across replicates by taking the mean. For each cell line separately, we corrected LFCs for gene copy number alteration using relative copy numbers provided by CCLE using the methodology described in Meyers et al. [2017].

### 2.4 List of paralog gene pairs

We downloaded human gene paralog pairs from the Protein ANalysis THrough Evolutionary Relationships (PANTHER) database [Mi et al., 2016] using the file ftp://ftp.pantherdb.org/ortholog/13.1/ RefGenomeOrthologs.tar.gz. Paralogs, defined as genes that diverged via a duplication event, were predicted using phylogenetic trees of protein-coding genes across 104 organisms. We only considered genes screened in the Achilles dataset, resulting in 74,070 paralog pairs.

## 3 Results

### 3.1 The impact of multiple on-target alignments on sgRNA depletion

We investigated the effects of multiple on-target alignments by looking at the relationship between sgRNA alignments and LFCs. We note that negative LFCs indicate a decrease in cell proliferation, and therefore larger negative LFCs indicate greater gene essentiality. For our analyses, we also corrected LFCs for copy number variation using the methodology described in Meyers et al. [2017]. In Figure 1a, we present the counts of guides stratified by the number of targets; the counts decay exponentially as the number of perfect alignments increases. 68,742 guides align uniquely to one target only, and 3,959 guides align to more than one target (multi-target guides), resulting in 2023 genes that are targeted by at least one multi-target guide. Interestingly, 86 sgRNAs intended to target genes did not align to any genomic location in GRCh38. The median LFC for these guides is positive (Figure 1b) and correlates with the 995 non-targeting control (NTC) guides included in the Avana library (*r* = 0.69), suggesting indeed that many of the DNA sequences corresponding to those 86 sgRNA sequences are likely to be absent in the genome.

**Figure 1.**
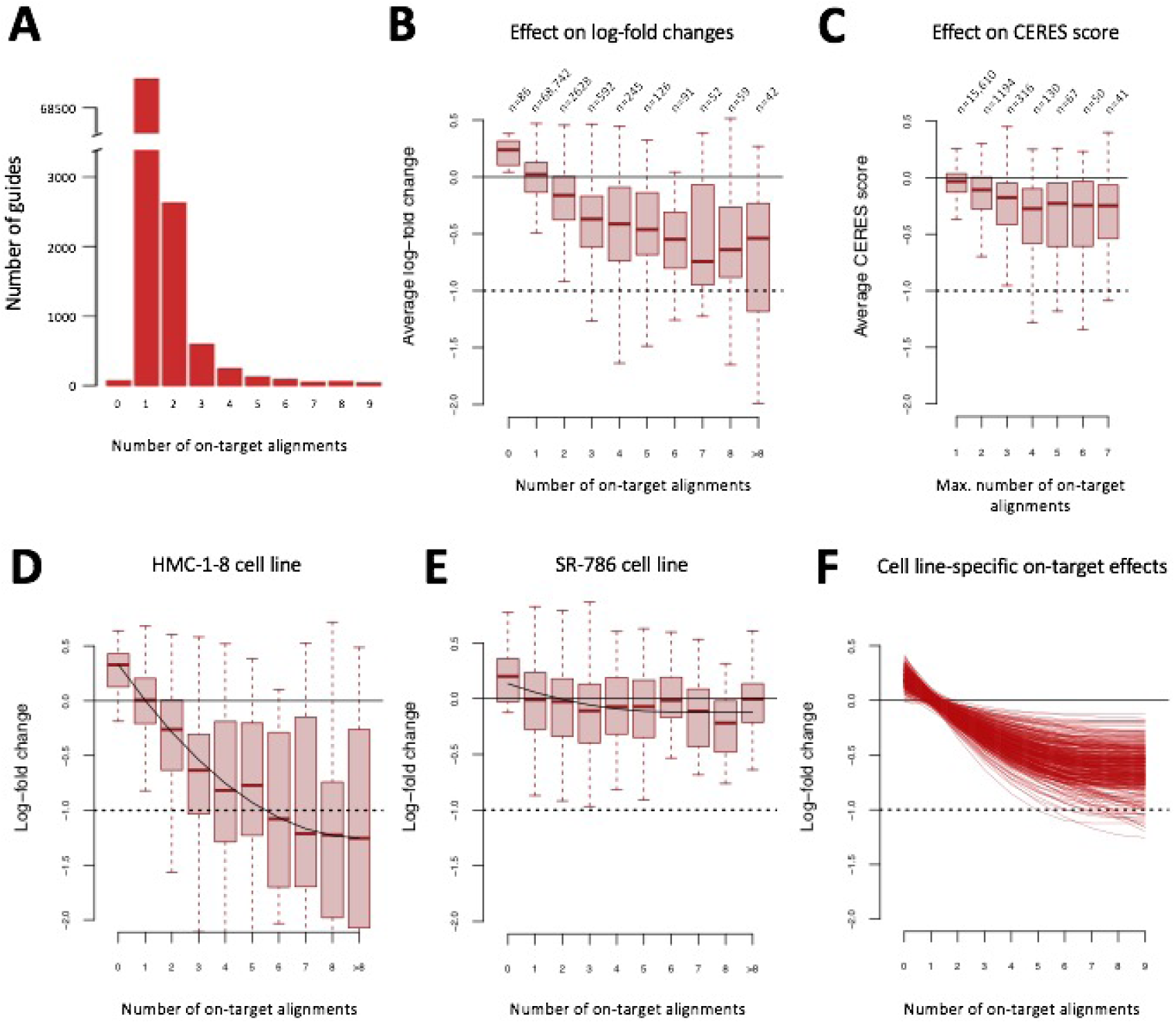
The impact of multiple on-targets on sgRNA log-fold changes. **(a)** Guide numbers as a function of the number of perfect alignments (on-target alignments). **(b)** Guide-level log-fold changes (LFCs) averaged across cell lines as a function of the number of on-targets. The number of on-targets was calculated as the number of perfect alignments between the reference genome and the 20-nt protospacer; we excluded guides with single-mismatch alignments to prevent the confounding effect of single-mismatch off-targets. **(c)** Combined effect of multiple on-targets on the CERES score. For each gene, we calculated the maximum number of targeted loci (x-axis) as the maximum number of perfect alignments for a guide designed to target that gene in the Avana library. **(d-e)** Effects of multiple on-target alignments on LFCs in the breast cancer cell line HMC-1-8 and in the lymphoma cell line SR-786. The solid lines represent second-degree polynomial fits (see Methods). **(f)** Fitted on-target activity score as a function of the number of on-target alignments; each fitted line is a separate cell line

To study the impact of multiple alignments on guide essentiality scores, we averaged LFCs across cell lines to get guide-specific average essentiality scores. We restricted our analysis to guides with no single-mismatch alignments to prevent confounding with off-target effects; off-target effects caused by mismatch tolerance are further discussed below. In Figure 1b, we report the distributions of the average LFCs as a function of the number of perfect alignments. The median LFC significantly decreases as a function of number of perfect alignments (p < 2.2 × 10^−16^, Jonckheere trend test [Jonckheere, 1954, Terpstra, 1952]). This is concordant with the hypothesis that a guide mapping to several DNA targets will introduce multiple DSBs and therefore will result in more cleavage toxicity, similar to the effect of copy number [Meyers et al., 2017]. The CERES algorithm described in Meyers et al. [2017] attempts to account for these multi-target effects by explicitly modeling multi-target gene knockouts in an additive fashion. Despite this implementation, we observed that the number of perfect alignments also affects the median CERES score (p < 2.2 × 10^−16^, Jonckheere trend test, Figure 1c).

We also found that cleavage toxicity induced by multi-target guides is cell line-specific, similar to the CN toxicity described in Meyers et al. [2017]; for instance, the on-target toxicity is quite profound for the breast cancer HMC-1-8 cell line (Figure 1d), whereas there is minimal to no effect in the lymphoma cell line SR-786 (Figure 1e). For both cell lines, the fitted curves were estimated using second-degree polynomials (see Methods). Fitted curves for all cell lines are presented in Figure 1f.

### 3.2 Guides co-targeting coding regions are enriched for paralogs

Next, we sought to understand whether or not the increased lethality associated with multi-target guides is attributable to cleavage toxicity only, or can also be a consequence of genetic interactions within the set of co-targeted genes. We focused on the 2628 guides in the Avana library that target exactly two genomic loci with perfect complementarity; we refer to these guides as “double-target” guides. A double-target guide can either target (a) one coding region and one non-coding region, or (b) two coding regions. For (a), the combined knockout effect is expected to be the sum of the gene-specific knockout effect and the cleavage toxicity effect induced by introducing DSBs at two genomic loci. For (b), the combined knockout effect is expected to be the sum of the double-gene knockout effect, with possibly a genetic interaction effect, and the cleavage toxicity effect induced by introducing DSBs at two genomic loci.

Using annotated exons from GENCODE (comprehensive gene annotation, human v28), we found that 2503 (95.2%) of the double-target guides have both targets located in coding regions. Since the annotated exons represent only 4.5% of the mappable genome, this represents a significant enrichment (exact binomial test, p < 2.2 × 10^−16^). We studied the effects of coding vs non-coding targets by averaging LFCs across cell lines for double-target guides. We excluded guides with single-mismatch off-targets, resulting in 1734 and 85. guides for coding and non-coding region secondary targets, respectively. The average LFCs are presented in Supplementary Figure S1. For both sets of guides, the median LFC is comparable and below 0 as a result of cleavage toxicity induced by introducing DSBs at two genomic loci. However, the number of guides with high activity (guides with LFC ≤ − 0.5) is significantly higher for the set of guides targeting two coding regions (OR = 4.32, p = 0.00094, Fisher’s exact test) than for the set of guides targeting only one coding region. This suggest that a guide disrupting two genes is likely to be more lethal than a guide targeting one coding region and one non-coding region.

For guides targeting two coding regions in the Avana library, we asked whether or not the two targets are related in terms of sequence similarity using gene paralogy as a proxy for gene similarity. Among the 2503 guides, 297 (11.9%) guides have their pair of targets annotated as being paralogs using the PANTHER database [Mi et al., 2016] (see Methods). In comparison, the 74,070 pairs of paralog genes annotated in the PANTHER database represent only∼ 0.02% of all possible pairs of genes screened in the Avana library. This significant enrichment for paralog genes (exact binomial test, p < 2.2 × 10^−16^) confirms that co-targeted genes often share high homology.

### 3.3 The interplay between multi-target guides and synthetic lethality: *MYL12A, MYL12B* and *MYL9*

In Meyers et al. [2017], guide-level LFCs are modeled as a combination of multiple on-target gene knockout effects. The model makes the assumption that gene knockout effects are additive: the growth phenotype resulting from double mutant cells is the same as the sum of the single-mutant growth phenotypes. While this assumption is likely valid for pairs of genes/loci with no genetic interaction, such as DSB effects in non-coding regions, this can lead to erroneous estimates of gene essentiality in case of synergistic or epistatic genetic interactions. Many pairs of paralogous genes are functionally, or at least partially, redundant, and synergistic effects have been observed for such pairs [Pérez-Pérez et al., 2009]. We therefore expected some of the sgRNAs co-targeting paralog genes to violate the additivity assumption and result in biased estimates of gene essentiality.

As an example, synthetic lethality, in which deficiencies in two (or more) genes is lethal while deficiency in either one is not, is a genetic interaction that cannot be captured through additive models. The myosin light chain 12A (*MYL12A*) and myosin light chain 12B (*MYL12B*) genes, two paralogous genes that are part of the myosin II complex, are both targeted by unique and common guides in the Avana library (see Figure 2a and Supplementary Table S1). Guides A1 and A2 map uniquely to *MYL12A* while guides B1 and B2 map uniquely to *MYL12B*. Guides B3 and B4 map to *MYL12B*, but also to the processed pseudogenes *MYL12BP1* and *MYL8P*. Finally, guide AB1 maps to both *MYL12A* and *MYL12B* (two genes), while AB2 maps to *MYL12A* and *MYL12B* in addition to *MYL12AP1, MYL12BP1, MYL12BP2* and *MYL8P* (six genes/pseudogenes).

**Figure 2.**
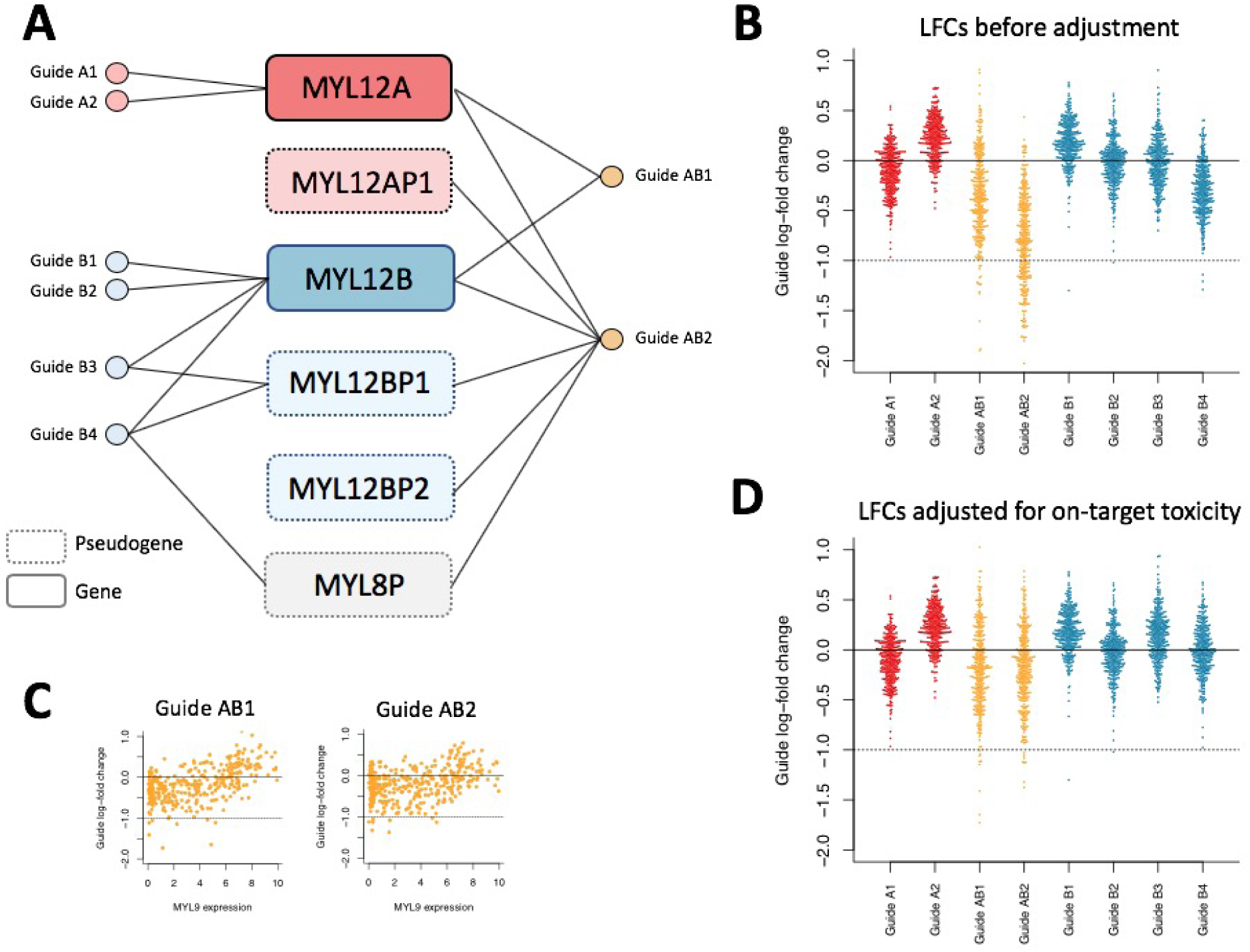
The interplay between multiple alignments and synthetic lethality in the Avana library. **(a)** Genomic mapping of the Avana guides targeting *MYL12A* and *MYL12B*. Guides A1 and A2 map uniquely to *MYL12A* and guides B1 and B2 map uniquely to *MYL12B*. Guides B3 and B4 map to *MYL12B*, but also to additional non-functional pseudogenes. Guides AB1 and AB2 map to both *MYL12A* and *MYL12B*. **(b)** Copy number-corrected LFCs for guides mapping to either *MYL12A* or *MYL12B*, or to both, in the Avana library, across 391 cell lines; each dot represents a cell line. **(c)** Relationship between guide-specific log-fold changes and *MYL9* expression for the two guides mapping to both *MYL12A* and *MYL12B*. **(d)** Same as (b), but after adjusting for cell-specific cleavage toxicity induced by multiple on-target alignments.

CN-corrected LFCs for these 8 guides are presented in Figure 2b. The LFC distributions for the isoform-specific guides (A1, A2, B1, B2 and B3) are approximately centered around 0. The guide targeting two pseudogenes (B4) appears to be more active; this is consistent with the toxicity effect observed in Figure 1b for guides targeting 3 loci. Interestingly, for guides targeting both *MYL12A* and *MYL12B*, the LFCs are shifted downwards with more variability, indicating that these two guides targeting both isoforms are substantially more toxic. For both guides, it appears that a subset of cells lines have a depletion score comparable to that of an essential gene (around −1). This suggests that the double knockout of *MYL12A* and *MYL12B* is potentially lethal for a subset of cell lines, suggesting context-dependent synthetic lethality between the two paralogs.

Using our estimation of cell-specific cleavage toxicity induced by multiple on-target alignments presented in Figure 1f, we can adjust LFCs for global on-target toxicity; adjusted LFCs are presented in Figure 2d. The scores within a guide category (A,B or AB) tend to be more similar after adjustment. It is also clear that additive on-target activity is not sufficient to fully explain greater activity of guides AB1 and AB2. This suggests that an additive model without genetic interaction does not capture the underlying biology and leads to erroneous estimates.

To formally test for a genetic interaction between *MYL12A* and *MYL12B*, we model the LFC *y*_*i*_ for guide *i* with the linear model

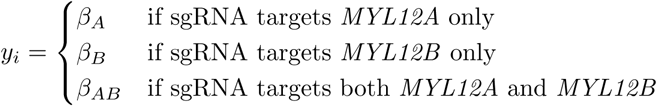

and test whether or not *β*_*A*_ + *β*_*B*_ = *β*_*AB*_. Using all cell lines to estimate the parameters, we obtained the isoform-specific knockout effects 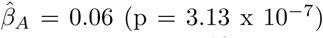 and 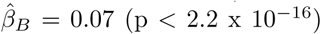, and the digenic knockout effect 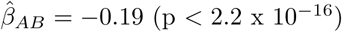. From the theory of linear modeling, we can obtain a t-statistic and its associated p-value to test the additivity hypothesis *β*_*A*_ + *β*_*B*_ − *β*_*AB*_ = 0. We obtained a significant test (p *<* 2.2 × 10^−16^) and conclude that the additivity hypothesis does not hold, therefore confirming a synergistic effect between *MYL12A* and *MYL12B*.

To investigate in which genomic context the genetic interaction between *MYL12A* and *MYL12B* appears to be maximal, we looked at the correlation between LFCs for Guides AB1 and AB2 and gene expression of 23,241 genes obtained from CCLE (see Methods). *MYL9* was the top correlate for both guides (Guide AB1: *r* = 0.534, p *<* 2.2 × 10^−16^; Guide AB2: *r* = 0.377, p = 1.6 × 10^−13^; see Figure 2c). None of the guides were associated with either *MYL12A* and *MYL12B* expression. This suggests that the pair of paralogs is more essential to cell survival in the absence of *MYL9* expression. Interestingly, both *MYL12A* and *MYL12B* are non-muscle regulatory light chains (RLCs) that are highly homologous to the RLC *MYL9*. The murine orthologs (*Myl12a, Myl12b* and *Myl9*) have been shown to be required to maintain the stability of myosin II and cellular integrity, and double knockdown of *Myl12a/Myl12b* using siRNA showed major alterations in cell structure that were not recapitulated by isoform-specific knockdowns [Park et al., 2011].

### 3.4 Additive models can lead to gene essentiality scores highly dependent on guide design

Besides the problem of genetic interactions, the use of additive models to estimate gene knockout effects from multi-target guides can lead to further problems. We present in this section two guide designs that are part of the Avana library that lead to CERES scores that need to be interpreted with caution because of a violation of the additivity assumption.

In Figure 3a, we illustrate the guide design for the Avana guides targeting the two genes *TMED7* or *TICAM2* as well as the readthrough *TMED7-TICAM2*. Guides 1-4 target both *TMED7* and *TMED7-TICAM2*, Guides 5-7 target both *TICAM2* and *TMED7-TICAM2*, while Guide 8 targets only *TICAM2*. Using the additive model posited in the CERES algorithm, gene scores for *TMED7, TICAM2* and *TMED7-TICAM2* can be solved using ordinary least squares (OLS). We note that in the CERES model, two additional guide-specific parameters are included in the model to capture a guide-specific activity score and offset; an iterative least squares approach is used to iteratively solve for guide-specific parameters and gene essentiality scores. These two location-scale parameters do not alter the interpretation of the gene essentiality scores derived from the additive model. Consequently, we omit them here for simplicity in order to focus on the guide-specific test of genetic interaction. For a specific cell line, let *y*_1_, *y*_2_, *…, y*_8_ denote the CN-corrected LFCs for Guides 1-8 respectively using the guide notation presented in Figure 3a. An additive model for the gene essentiality scores *β*_TMED7_, *β*_TICAM2_ and *β*_Fusion_ can be represented using the following system of linear equations:

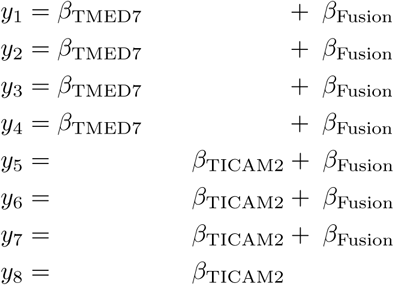

**Figure 3.**
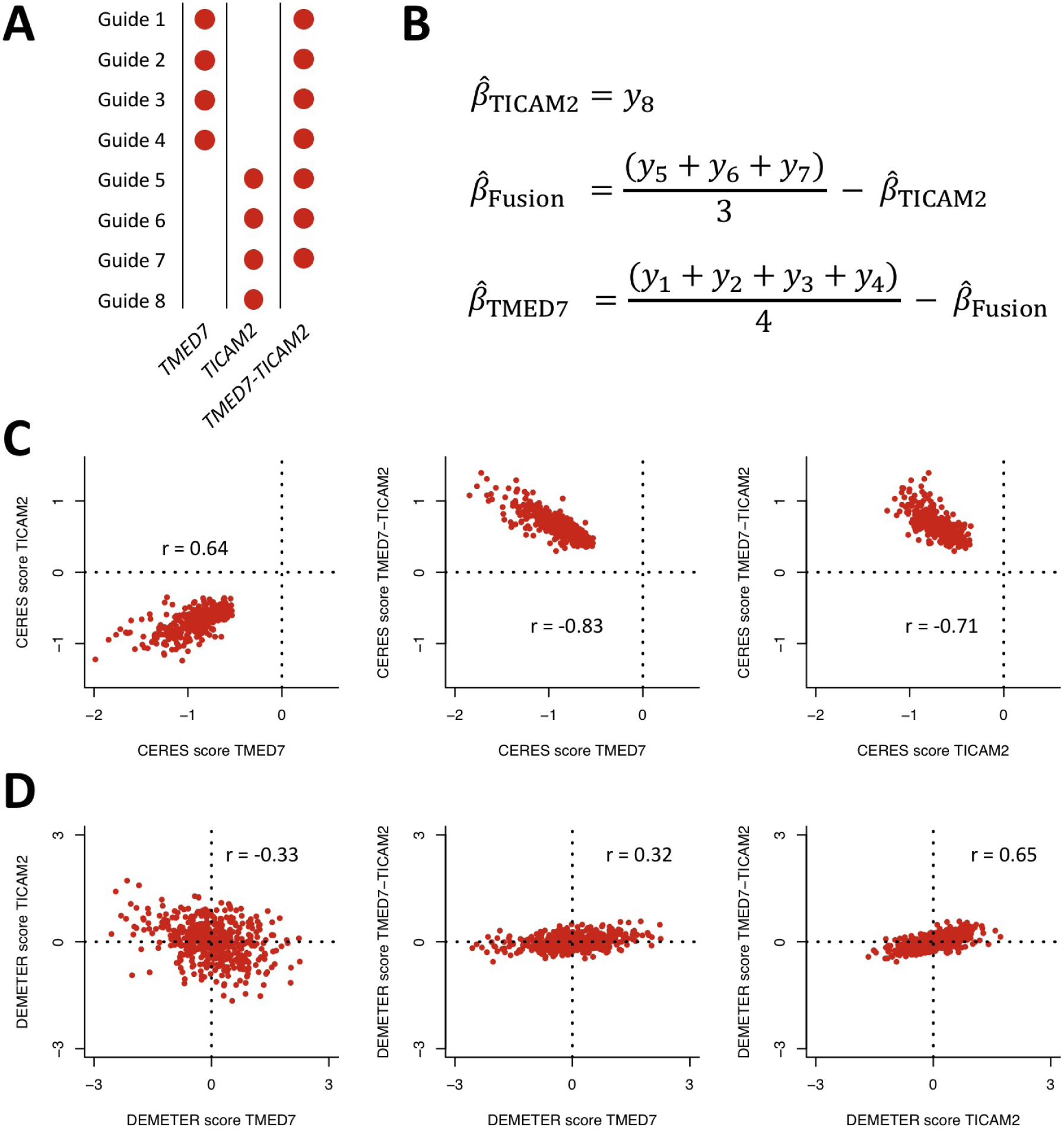
Analysis of the genetic knockout additivity constraints for *TMED7, TICAM2* and the *TMED7-TICAM2* readthrough. **(a)** Design for guides targeting *TMED7, TICAM2* and the readthrough *TMED7-TICAM2* in the Avana library. **(b)** Cell line-specific gene score solutions for an additive model using the Avana guides targeting *TMED7, TICAM2* and the readthrough *TMED7-TICAM2*. *y*_*i*_ denotes the LFC for guide *i* and *β*_*Fusion*_ denotes the gene score for the readthrough *TMED7-TICAM2*. Explicit solutions are derived using ordinary least squares (OLS). **(c)** Pairwise scatterplots for the CERES scores for *TMED7, TICAM2* and the readthrough *TMED7-TICAM2*. **(d)** Pairwise scatterplots for the DEMETER scores (RNAi) for *TMED7, TICAM2* and the readthrough *TMED7-TICAM2*.

The OLS estimates for gene essentiality scores can be written as

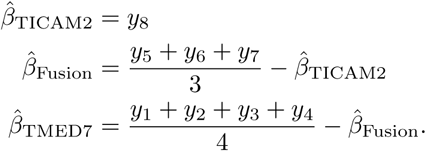

The gene essentiality score for *TICAM2* depends entirely on the LFC of one guide (Guide 8) and is therefore highly sensitive to outliers and off-target effects. The essentiality scores for both *TMED7* and *TMED7-TICAM2* depend on that of 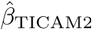, making the scores inter-dependent and creating a correlation structure highly dependent on guide design, and not necessarily representative of true on-target effects. In the Achilles dataset, we found that the CERES scores for these three genes are highly correlated with each other (Figure 3c) and consistent with the linear dependencies introduced by the OLS solution shown in Figure 3b and discussed above. To examine whether or not the CERES estimates for these three genes are biased, we studied the DEMETER scores from the Achilles RNAi dataset [Tsherniak et al., 2017], presented in Figure 3d. Interestingly, the sign of the correlations between the three genes are reversed, and the three genes are broadly estimated as being non-essential genes (DEMETER score ≥ −2). This disagrees with the CERES scores of *TMED7* and *TICAM2*, which are centered at −1 and therefore are comparable to CERES scores of essential genes.

The influence of guide design on correlations between essentiality scores is not limited to the triplet *TMED7* / *TICAM2* /*TMED7-TICAM2*. The Avana library targets 36 readthrough genes, and all of them are targeted by guides that are not unique to the readthrough but also target the pairs of individual genes. The correlations of the CERES scores between two genes composing a readthrough have a distribution with two modes bounded away from 0 (Supplementary Figure S2, red line) which behave differently than pairs of genes chosen at random (grey line). This suggests that the guide design introduces spurious correlations between gene essentiality scores for most genes targeted in a readthrough.

Spurious correlations can also happen with pairs of genes targeted by multi-target guides, such as the pair *EIF3C* /*EIF3CL*. These two highly homologous genes are part of the eukaryotic translation initiation factor 3 complex (eIF3) and therefore are expected to be essential for cell growth. In the Avana library, five guides target both *EIF3C* and *EIF3CL*, and one additional guide targets *EIF3CL* only. All six guides have LFCs centered around −1, indicating gene essentiality, yet the mean CERES scores for *EIF3CL* and *EIF3C* are respectively −1.15 and 0.20, suggesting that *EIF3CL* is broadly essential and *EIF3C* broadly non-essential. This contradicts the findings of Hart et al. [2014], which report *EIF3C* as a pan-cancer essential gene. This is a consequence of the knockout additivity assumption; the fact that the LFCs for the 5 guides targeting both paralogs is similar to the LFC of the guide targeting only *EIF3CL* leads to an estimate of the *EIF3C* CERES score close to 0. However, assuming both genes are broadly essential, it seems equally plausible that the double knockout does not make the cells die more in comparison to single knockout, especially since the single knockouts already induces a strong cell killing. The similar yet non-redundant function of the two genes violates the additivity assumption and leads to an incorrect estimate of the gene essentiality score for *EIF3C*. These two examples suggest that multi-targeting can lead to guide design-dependent codependencies and misleading biases that have to be interpreted with caution in downstream applications such as identifying gene networks and cancer cell dependencies.

### 3.5 The impact of mismatch tolerance on sgRNA depletion

We now focus on characterizing off-target effects caused by mismatch tolerance between the sgRNA’s protospacer sequence and the genomic DNA. In Figure 4a, we plot the guide counts distribution as a function of the number of single-mismatch alignments; a single-mismatch alignment is defined as a one-nucleotide mismatch in the sgRNA-DNA pairing. To delineate cleavage toxicity caused by multiple-target alignments (on-target effects) from off-target effects due to mismatch tolerance, we stratified guides by the number of single-mismatch alignments and the number of perfect alignments (Figure 4b). For a fixed number of perfect alignments, additional single-mismatch alignments significantly decrease LFCs (multiple linear regression, p *<* 2.2 × 10^−16^). This confirms that in addition to perfect alignments, single-mismatch off-targets contribute independently and additively to the decrease of cell viability. Similar to the cleavage toxicity associated with multiple-target guides, we found that off-target toxicity is also cell line-specific (Figure 4c). Moreover, cell line-specific off-target toxicity correlates with cell line-specific on-target toxicity (*r* = 0.69, p *<* 2.2 × 10^−16^); we used the fitted on-target and off-target effects for 4 perfect and single-mismatch alignments, respectively, as measures of on-target and off-target toxicity. The correlation suggests that both on-target and off-target cleavage toxicity are related effects that show specificity for cell lines but not guides, since the sets of guides used to estimate both effects are broadly different (see Methods). Furthermore, cleavage toxicity anti-correlates with the cell line-specific Cas9 activity score described in Aguirre et al. [2016] (*r* = − 0.48, p *<* 2.2 × 10^−16^, Figure 4e), and strongly anti-correlates with the median LFC for non-targeting controls (*r* = 0.67, p *<* 2.2 × 10^−16^).

**Figure 4.**
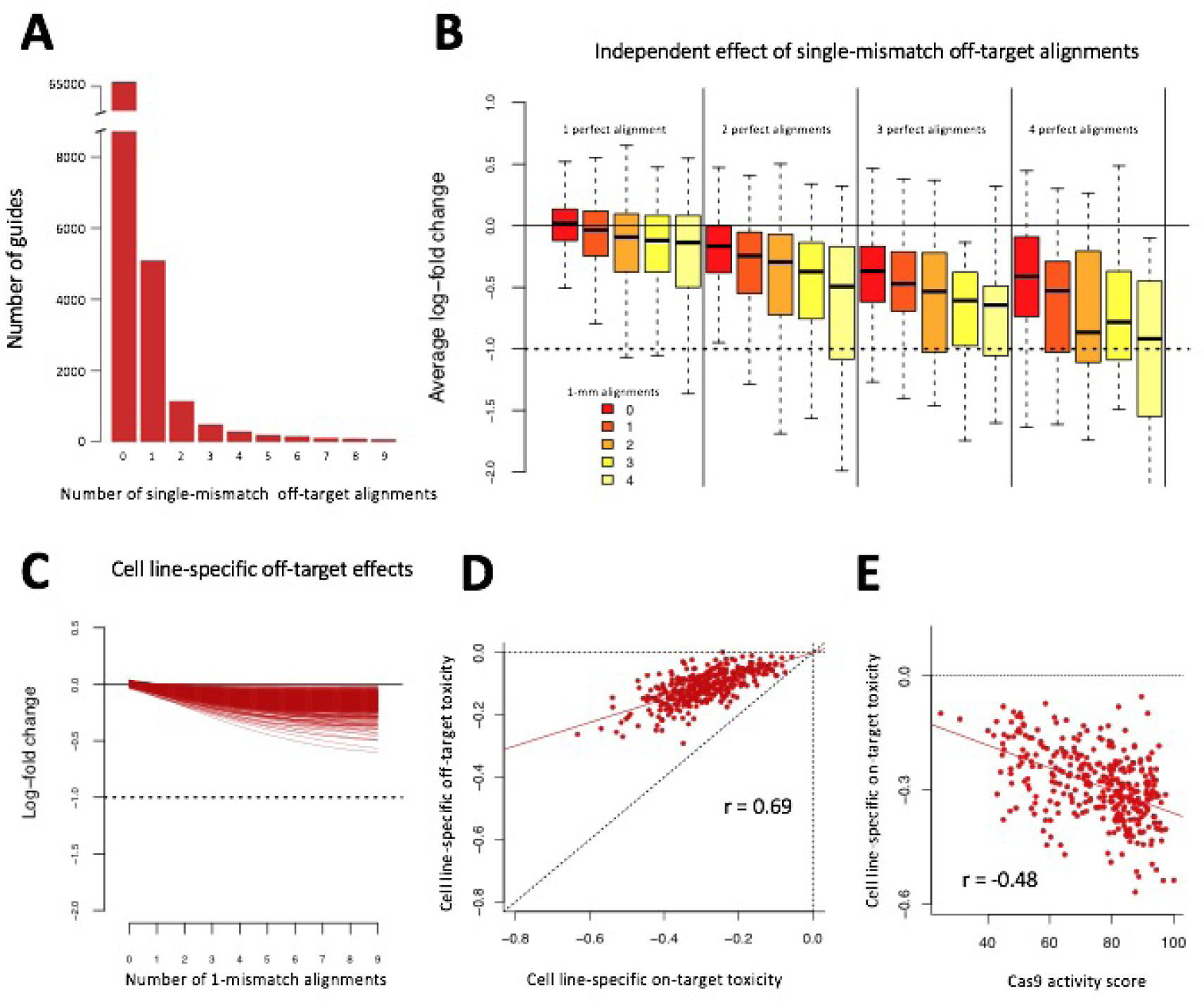
The impact of off-targets on sgRNA log-fold changes. **(a)** Guide counts as a function of the number of single-mismatch alignments. **(b)** Combined effect of multiple perfect alignments (on-targets) and single-mismatch (1-mm) alignments on LFCs. **(c)** Fitted off-target effects caused by single-mismatch alignments. Each fitted line is a separate cell line. **(d)** Relationship between cell line-specific on-target toxicity and cell line-specific off-target toxicity. On-target and off-target toxicity were measured as the average LFC for guides with 4 perfect alignments and 4 single-mismatch alignments, respectively. **(e)** Relationship between on-target toxicity and Cas9 activity score.

We also studied the relationship between mismatch position in the protospacer and mismatch tolerance across cell lines. To prevent the number of alignments from confounding the analysis, we only considered guides with one perfect alignment and one single-mismatch alignment. In Figure 5a, we show the distributions of the LFCs as a function of single-mismatch position within the protospacer. The effect of a single mismatch is more pronounced for mismatches far away from the PAM site (PAM-distal region) as opposed to PAM-proximal nucleotides, sometimes referred to as the seed region. Mismatch tolerance appears to be maximal at the 20th position. This suggests that guides should be carefully designed to avoid mismatch at that position.

**Figure 5.**
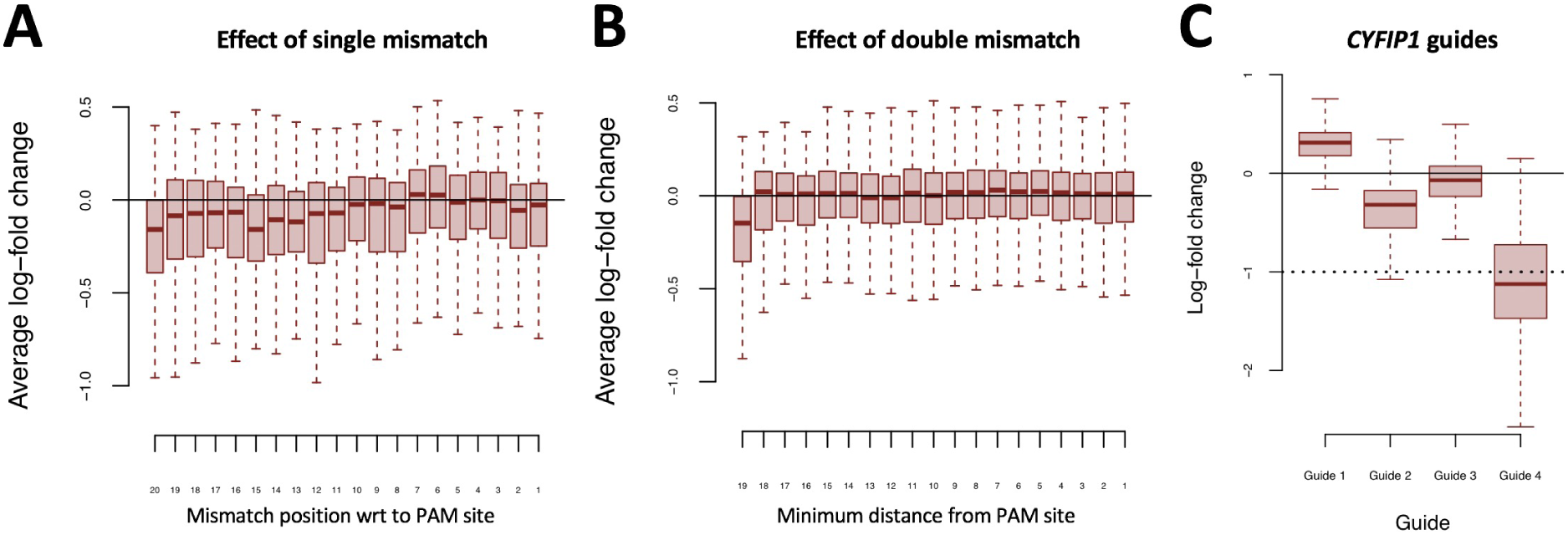
Position-specific mismatch tolerance of the protospacer. **(a)** Effect of a single mismatch between sgRNA protospacer and target DNA as a function of position within the protospacer. The PAM site starts at position 0. **(b)** Effect of a double mismatch between protospacer and reference genome as a function of the most PAM-proximal mismatch position. **(c)** Log-fold changes across cell lines for the 4 guides targeting *CYFIP1*. The protospacer of Guide 4 aligns to 10 genomic loci when allowing a mismatch at both position 20 and position 19.

We extended our analysis to double-mismatch (sgRNA-DNA mismatch at two nucleotides) tolerance. Because the number of all possible combinations of two-nucleotide mismatches in the 20-nt protospacer is large compared to the number of available guides with mismatches in the Avana library, we confined our analysis to pairs of mismatches for which the two discordant bases are located within a specified distance from the PAM site. The distributions of guide depletion are shown in Figure 5b. Again, to prevent the number of alignments from confounding the analysis, we only considered guides with one double-mismatch alignment. One can observe an apparent off-target effect for a double mismatch that occurs at the two most PAM-distal positions. To illustrate the impact of double mismatch on guide essentiality, we consider guides targeting the gene *CYFIP1*. The LFCs across cell lines are illustrated in Figure 5c. All four guides have only *CYFIP1* as an on-target alignment, and do not have any single-mismatch alignments. Guide 1 has one double-mismatch alignment (positions 3 and 12); guides 2 and 3 do not have any double-mismatch alignment. Guide 4 aligns to 10 different genomic loci with a double-mismatch at positions 19 and 20, resulting in a substantial decrease in the LFC. This biases the CERES score for *CYFIP1* towards essentiality (CERES score of − 0.41), while excluding guide 4 results in a score centered around 0 (non-essentiality).

### 3.6 Mismatch-tolerant guides can confound phenotypes: *SOX9* and *SOX10*

The CERES model does not account for off-target effects caused by single-mismatch and double-mismatch tolerance, and this can lead to erroneous conclusions when both on- and off-targets are part of the same gene family, for instance genes within the SOX family. Importantly, we found that among the 4705 genes that have at least one guide with a single-mismatch alignment, 3197 (68%) such genes have at least one single-mismatch alignment located in the exon of another gene. This increases the likelihood of false positive effects caused by single-mismatch alignments. The consequences are potentially variable across cell lines and may depend on which alternative family member is affected, and whether it is expressed or not in a given subset of cell lines. We illustrate this by analyzing the CERES score and guide LFCs for the *SOX9* gene encoding the transcription factor SOX-9.

By looking at the relationship between the CERES score for *SOX9* and its expression (Figure 6a), we noticed quite a few cell lines with very low expression of *SOX9* that exhibit a clear and contradictory dependency on *SOX9* (CERES score close to −1); most of these cell lines are melanoma cell lines. While all of the protospacer sequences for the guides targeting *SOX9* match perfectly to only *SOX9*, an analysis of single-mismatch alignments revealed that 3 out of 4 such guides also align to *SOX10*. By plotting the *SOX9* CERES score against *SOX10* expression, it becomes clear that *SOX9*-dependent cell lines with low expression of *SOX9* are those that highly express *SOX10*. ((Figure 6b).

**Figure 6.**
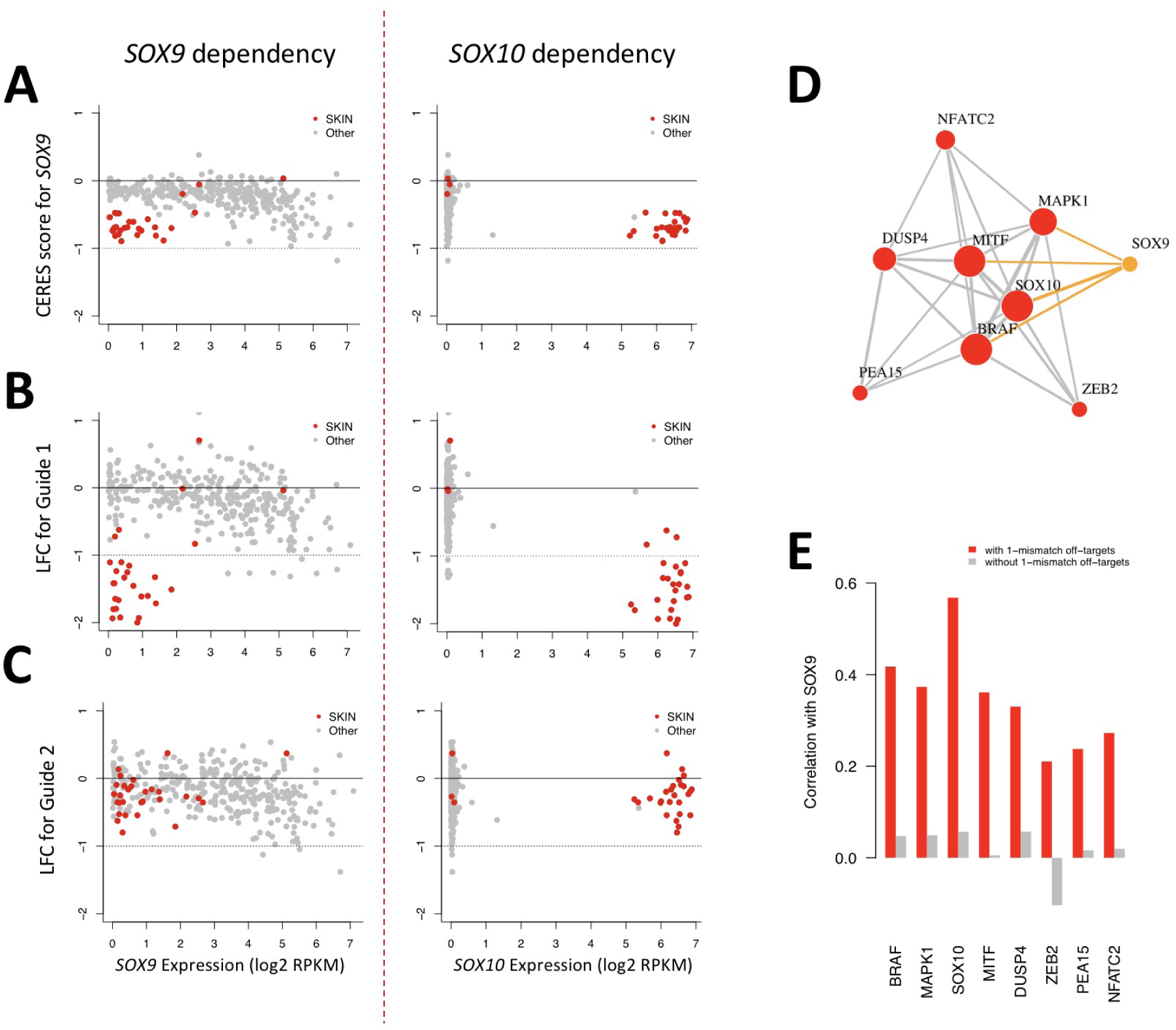
The confounding effect of single-mismatch sgRNA-DNA tolerance. **(a)** On the left: CERES score for *SOX9* plotted against *SOX9* expression; each dot is a cell line, and skin cell lines are plotted in red. On the right: CERES score for *SOX9* plotted against *SOX10* expression. **(b)** Same as (a), but with the CN-corrected log-fold change for a guide targeting *SOX9* with a single-mismatch alignment to *SOX10*. **(c)** Same as (b), but for a guide without a single-mismatch alignment to *SOX10*. **(d)** *BRAF*-associated coessentiality cluster using between-gene CERES Pearson correlations. A correlation cutoff of *r* = 0.35 was chosen to draw edges. Edges associated with *SOX9* (orange) disappear when not including *SOX9*-targeting guides with 1-mismatch to *SOX10*. **(e)** Pearson correlations between the *SOX9* CERES score and genes that are part of *BRAF*-associated cluster, before and after removal of guides with 1-mismatch *SOX10* off-target effects.

Analyzing guide-level LFCs reveal that indeed, for a guide with a single-mismatch alignment to *SOX10* (Guide 1), there is a strong dropout in cell lines highly expressing *SOX10* (Figure 6b). Conversely, based on a guide with no single-mismatch alignment to *SOX10* (Guide 2), cell lines expressing *SOX10* are not dependent on *SOX9* (Figure 6c).

Without analyzing single-mismatch alignments, one would have erroneously concluded that most of the melanoma cell lines in the Achilles dataset have a *SOX9*, rather than a *SOX10*, dependency. This illustrates how single-mismatch off-target effects can confound the analysis of cancer vulnerabilities, and also how it can be challenging to detect those off-target effects when they occur only in a small subset of cell lines, such as the melanoma cell lines in this case.

Recently, a coessentiality network has been derived from correlating fitness profiles across CRISPR knockout screens datasets, with the Achilles dataset being the most represented dataset [Kim et al., 2018]. The authors found a coessentiality network cluster that is highly specific to *BRAF*-mutated melanoma cell lines and contains elements of the MAP kinase pathway (*MAP2K1, MAPK1* and *DUSP4*) as well as *SOX9* and *SOX10*. Using the CERES scores from the Achilles dataset, we were able to recreate the cluster entirely by taking the 9 top genes correlated with the *BRAF* CERES score: *BRAF, MITF, MAPK1, PEA15, NFATC2, ZEB21, DUSP4, SOX9, SOX10* (Figure 6d). The correlation between the CERES score for *BRAF* and the CERES score for *SOX9* is high (*r* = 0.42). We recalculated an essentiality score for *SOX9* after removing guides with 1-mismatch alignment to *SOX10*. As a result, *SOX9* no longer associates with the *BRAF* cluster (Figure 6e).

This is not an isolated case. By looking at the correlation between the CERES score and self-expression of the gene, we found that many of the top anti-correlated genes, which are overrepresented by transcription and lineage factors, have guides with single-mismatch alignments to an alternative member within their gene family: *GATA2* /*GATA3, SOX1* /*SOX2, DOCK10/DOCK11, UBB* /*UBC, PAX3* /*PAX7, TEAD2* /*TEAD3*, to name a few. We provide a table of the top 200 self-anti-correlated genes together with on-target and off-target alignments in the Supplementary File 2.

## 4 Discussion

In this work, we first analyzed cleavage toxicity associated with guides targeting more than one genomic locus with complete complementarity. Using LFCs from the Achilles dataset, across 342 cell lines and more than 75k sgRNAs (Avana library), we found that guide depletion increases as a function of the number of targeted loci in the genome. We observed this not only for perfect alignments between the sgRNA protospacer and genomic DNA, but also for single-mismatch tolerant alignments to a lesser extent. A single-mismatch that occurs in the 10 most PAM-distal nucleotides results in more severe toxicity, and a double-mismatch that occurs only in the 2 most PAM-distal protospacer positions results in greater toxicity. These biases have been reported before [Chen et al., 2018, Doench et al., 2016, 2014, Hsu et al., 2013, Xu et al., 2015, Fu et al., 2013, Anderson et al., 2015, Morgens et al., 2017, Tsai et al., 2015, Tsai and Joung, 2016, O’Geen et al., 2015, Lin et al., 2014, Kim et al., 2016, 2015], but were only estimated using a few cell lines. Here, we could robustly estimate these effects across hundreds of cell lines, and found that cleavage toxicity associated with promiscuous guides substantially depends on the cell line model, similar to the copy number bias detailed in Meyers et al. [2017]. The large number of cell lines and targeted genes made it possible to robustly generalize our findings across lineages and genetic contexts, despite the low number of guides per gene (average of 4 guides per gene in the Avana library).

The CERES algorithm presented in Meyers et al. [2017] implements a cell line-specific CN correction of the depletion scores, but also attempts to correct for multiple on-target effects by decomposing guide-specific LFCs as a sum of knockout effects. While this is valid for genes and genomic targets that do not interact with each other, such as DSBs introduced in non-coding DNA, the strict phenotypic additivity assumption often does not hold because of more complex genetic interactions. Two knockouts are considered to be strictly additive if the effect of the digenic knockout is the sum of the effects of the single knockout; strict additivity of cell fitness effects is rare, with most genes exhibiting some level of positive and negative genetic interaction which is substantially more frequent among essential genes [Costanzo et al., 2016]. Genetic interaction effects are also enhanced for guides targeting highly homologous genes that are more likely to function together in the same biological process or have some level of functional redundancy.

Synthetic lethality, for instance, is a type of genetic interaction in which double mutant cells do not survive, while single mutant cells continue to proliferate, perhaps at a slower rate. Pairs of synthetically lethal genes have been utilized to identify therapeutic targets: *BRG1-BRM* : [Hoffman et al., 2014, Januario et al., 2017], *ENO1-ENO2* [Muller et al., 2012], *ME2-ME3* [Dey et al., 2017], *TP53-POLR2A* [Liu et al., 2015] BRCA1/2-PARP[Farmer et al., 2005]. Using guides targeting both *MYL12A* and *MYL12B*, two regulatory light chains (RLCs) essential to the Myosin II complex, we showed that there exists a subset of cell lines for which the pair of RLCs is synthetically lethal, violating the assumption of additivity and culminating in biased CERES scores for both *MYL12A* and *MYL12B*.

We also showed that the additive model can create false inter-dependencies between genes in the presence of multi-target guides. Indeed, in the presence of a non-linear effect between a multigenic knockout and individual knockouts, the estimated gene knockout effects are markedly correlated with each other in patterns determined by the library guide design used in the experiment. We used the CERES scores estimated for *TICAM2, TMED7* and the readthrough *TMED7-TICAM2* to illustrate misleading CERES score correlations that are dependent on the single guide targeting *TICAM2* only.

Orthogonal to the problem of modeling multiple on-target knockout effects using an additive model, we also found that a single-mismatch in sgRNA-DNA alignments can confound phenotypic readouts because of off-target effects. We found that according to the CERES score, the transcription factor *SOX9* is essential in melanoma cell lines, despite the fact these cell lines lack expression of *SOX9* but highly express *SOX10*. We showed this was apparently caused by guides with a single-mismatch alignment to *SOX10*, causing off-target effects that confound the growth phenotype for *SOX9* non-expressing cell lines. Downstream consequences of such confounding was exemplified in a recent publication [Kim et al., 2018], in which the authors inferred a coessentiality network using the Achilles dataset and reported that *SOX9* is part of a gene cluster highly specific to *BRAF*-mutated melanoma cell lines. We showed that removing single-mismatch tolerant guides from the analysis removes *SOX9* membership in the cluster.

These observations suggest that multi-target guides as well as mismatch-tolerant guides can lead to false positives and biased essentiality scores. When interpreting essentiality score for a given gene, one should also investigate guide-level LFCs to detect abnormalities, such as guides with multiple on-target and off-target single-mismatch alignments. We provide in the Supplementary material a gene-level table summarizing the number of on-target and off-target alignments for the Avana library to help readers with flagging potentially problematic genes.

Similar to the CN correction algorithm implemented in Meyers et al. [2017], one could attempt to systematically correct for multi-target and off-target toxicity by removing the observed effects for each cell line separately. However, because of genetic interactions, correcting depletion scores for multi-target guides is not straightforward. As opposed to cleavage toxicity induced by differential CN, the increased activity observed in multi-target guides depends on the set of targeted genes; these genes can interact with each other in a cell line-specific manner, excluding the possibility of fitting a global genetic interaction correction model across cell lines. An alternative solution is to remove guides that do not map uniquely to the genome when calculating a gene-level essentiality score; 383 genes would have to be excluded in the Avana library because of the absence of uniquely-targeting guides for these genes. On the other hand, multi-target guides could be still further analyzed separately, since such guides can be informative about multigenic knockout, such as co-targeting known paralogs. These guides should be annotated separately, and may potentially be further utilized to help design a library of guides co-targeting paralogs.

## Abbreviations

CCLE: Cancer cell line encyclopedia
CRISPR: Clustered Regularly Interspaced Short Palindromic Repeats
DSB: DNA double-strand break
NSCLC: Non-small cell lung cancer
OLS: ordinary least squares
PAM: protospacer adjacent motif/
RLC: regulatory light chain
RNAi: RNA interference
sgRNA: single guide RNA
shRNA: short hairpin RNA siRNA: short interfering RNA.

## Authors contributions

JPF conceived and designed the study and performed the main computational analyses. PMH helped with writing and implementing software to process and visualize the Achilles data. JPF wrote the manuscript. JPF, MRC, SEM and WFF revised the manuscript with assistance from other authors. MRC, SEM, WFF and KEG helped with the biological interpretation of the results. All authors read and approved the final manuscript.

### Acknowledgements

We thank Bob Yauch, Matt Chang, Christiaan Klijn, Marc Hafner, Russell Bainer and Milena Dürrbaum for scientific discussions and technical assistance.

## Supplementary files

`supp table 1.csv`: genomic alignments to GRCh38 for the Avana guide sequences.

`supp table 2.csv`: top 200 self-correlated genes and single-mismatch alignments

`supp table 3.csv`: gene-level summary table for multiple on-target and off-target alignments

## Supplementary Tables and Figures

**Supplementary Figure S1.**
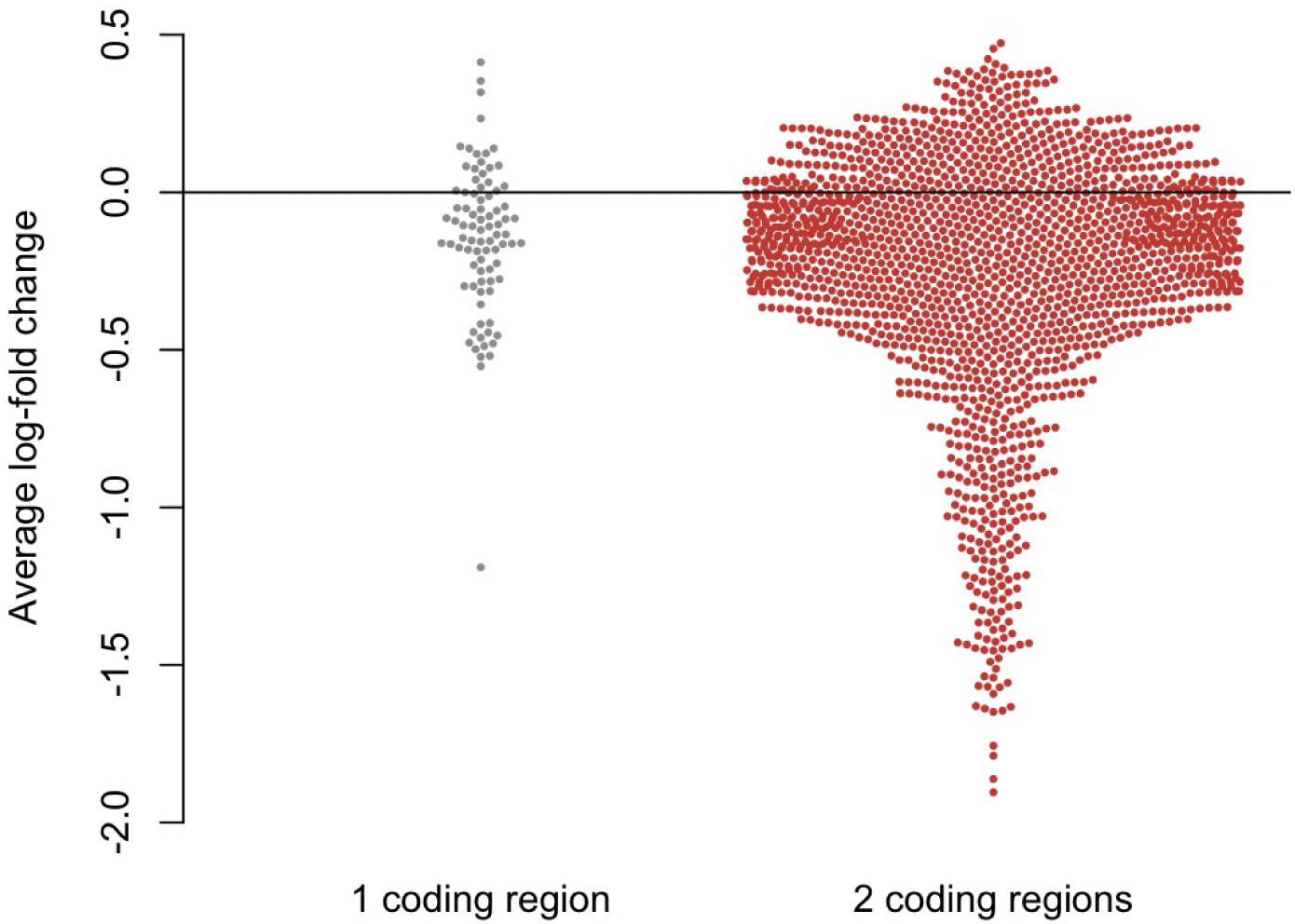
Log-fold changes for double-target guides. Log-fold changes averaged across cell lines for guides mapping to two genomic loci (double-target guides). They grey dots represent double-guides for which only one of the two targets is located in a coding region, and they red dots represent double-target guides for which both targets are located in coding regions.

**Supplementary Figure S2.**
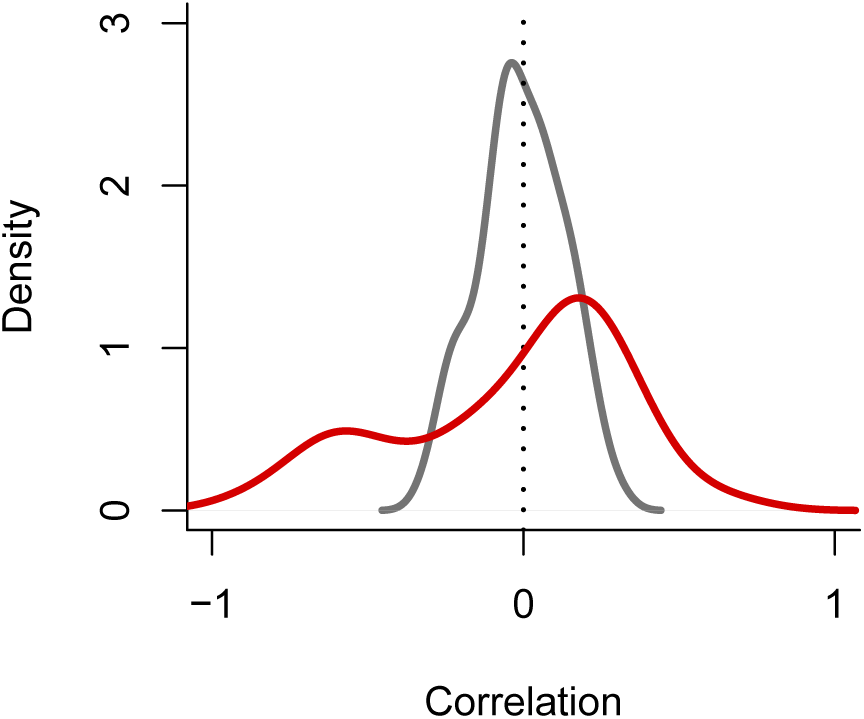
The effect of multi-target guides in readthrough genes. The Avana library contains 33 pairs of genes that can form a readthrough gene. The multi-target guides used to knock out individual genes as well as and the gene readthrough create inter-dependencies in the CERES scores. Red line: density of the pairwise correlations between the CERES score of genes for which their readthrough is also targeted. Grey line: pairwise correlations of random pairs of genes.

**Supplementary Table S1.**
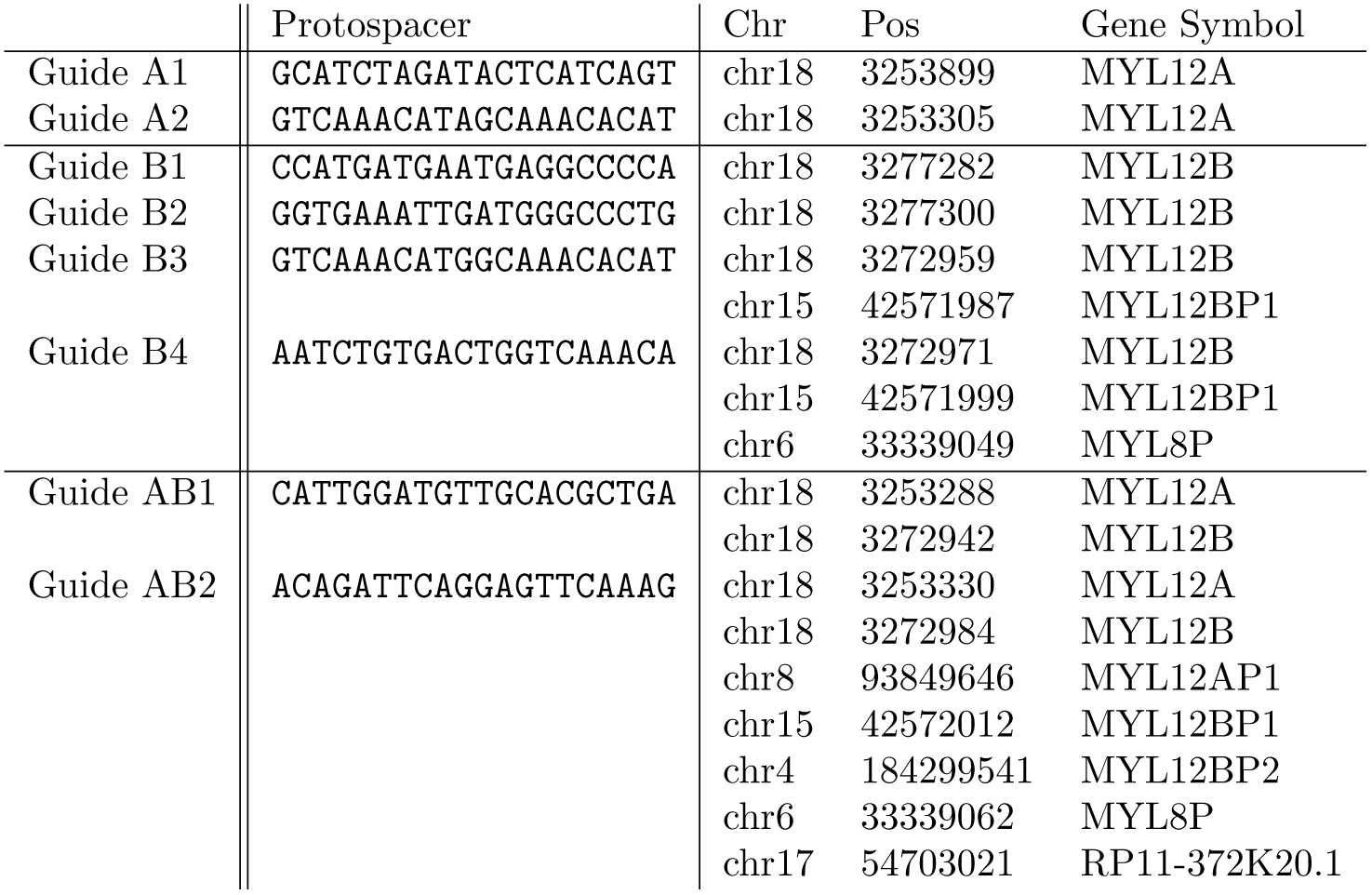
Genomic alignments for MYL12A and MYL12B guides in the Avana library.

## References

Andrew J Aguirre, Robin M Meyers, Barbara A Weir, Francisca Vazquez, Cheng-Zhong Zhang, Uri Ben-David, April Cook, Gavin Ha, William F Harrington, Mihir B Doshi, et al. Genomic copy number dictates a gene-independent cell response to crispr/cas9 targeting. Cancer discovery, 6(8):914–929, 2016.

Emily M Anderson, Amanda Haupt, John A Schiel, Eldon Chou, Hidevaldo B Machado, Zˇaklina Strezoska, Steve Lenger, Shawn McClelland, Amanda Birmingham, Annaleen Vermeulen, et al. Systematic analysis of crispr–cas9 mismatch tolerance reveals low levels of off-target activity. Journal of biotechnology, 211: 56–65, 2015.

Chen-Hao Chen, Tengfei Xiao, Han Xu, Peng Jiang, Clifford A Meyer, Wei Li, Myles Brown, and X Shirley Liu. Improved design and analysis of crispr knockout screens. Bioinformatics, 2018.

Michael Costanzo, Benjamin VanderSluis, Elizabeth N Koch, Anastasia Baryshnikova, Carles Pons, Guihong Tan, Wen Wang, Matej Usaj, Julia Hanchard, Susan D Lee, et al. A global genetic interaction network maps a wiring diagram of cellular function. Science, 353(6306):aaf1420, 2016.

Glenn S Cowley, Barbara A Weir, Francisca Vazquez, Pablo Tamayo, Justine A Scott, Scott Rusin, Alexandra East-Seletsky, Levi D Ali, William FJ Gerath, Sarah E Pantel, et al. Parallel genome-scale loss of function screens in 216 cancer cell lines for the identification of context-specific genetic dependencies. Scientific data, 1:140035, 2014.

Prasenjit Dey, Joelle Baddour, Florian Muller, Chia Chin Wu, Huamin Wang, Wen-Ting Liao, Zangdao Lan, Alina Chen, Tony Gutschner, Yaan Kang, et al. Genomic deletion of malic enzyme 2 confers collateral lethality in pancreatic cancer. Nature, 542(7639):119, 2017.

John G Doench, Ella Hartenian, Daniel B Graham, Zuzana Tothova, Mudra Hegde, Ian Smith, Meagan Sullender, Benjamin L Ebert, Ramnik J Xavier, and David E Root. Rational design of highly active sgrnas for crispr-cas9–mediated gene inactivation. Nature biotechnology, 32(12):1262, 2014.

John G Doench, Nicolo Fusi, Meagan Sullender, Mudra Hegde, Emma W Vaimberg, Katherine F Donovan, Ian Smith, Zuzana Tothova, Craig Wilen, Robert Orchard, et al. Optimized sgrna design to maximize activity and minimize off-target effects of crispr-cas9. Nature biotechnology, 34(2):184, 2016.

Hannah Farmer, Nuala McCabe, Christopher J Lord, Andrew NJ Tutt, Damian A Johnson, Tobias B Richardson, Manuela Santarosa, Krystyna J Dillon, Ian Hickson, Charlotte Knights, et al. Targeting the dna repair defect in brca mutant cells as a therapeutic strategy. Nature, 434(7035):917, 2005.

Yanfang Fu, Jennifer A Foden, Cyd Khayter, Morgan L Maeder, Deepak Reyon, J Keith Joung, and Jeffry D Sander. High-frequency offtarget mutagenesis induced by crispr-cas nucleases in human cells. Nature biotechnology, 31(9):822, 2013.

Jennifer Harrow, Adam Frankish, Jose M Gonzalez, Electra Tapanari, Mark Diekhans, Felix Kokocinski, Bronwen L Aken, Daniel Barrell, Amonida Zadissa, Stephen Searle, et al. Gencode: the reference human genome annotation for the encode project. Genome research, 22(9):1760–1774, 2012.

Traver Hart, Kevin R Brown, Fabrice Sircoulomb, Robert Rottapel, and Jason Moffat. Measuring error rates in genomic perturbation screens: gold standards for human functional genomics. Molecular systems biology, 10(7):733, 2014.

Gregory R Hoffman, Rami Rahal, Frank Buxton, Kay Xiang, Gregory McAllister, Elizabeth Frias, Linda Bagdasarian, Janina Huber, Alicia Lindeman, Dongshu Chen, et al. Functional epigenetics approach identifies brm/smarca2 as a critical synthetic lethal target in brg1-deficient cancers. Proceedings of the National Academy of Sciences, 111(8):3128–3133, 2014.

Patrick D Hsu, David A Scott, Joshua A Weinstein, F Ann Ran, Silvana Konermann, Vineeta Agarwala, Yinqing Li, Eli J Fine, Xuebing Wu, Ophir Shalem, et al. Dna targeting specificity of rna-guided cas9 nucleases. Nature biotechnology, 31(9):827, 2013.

Thomas Januario, Xiaofen Ye, Russell Bainer, Bruno Alicke, Tunde Smith, Benjamin Haley, Zora Modrusan, Stephen Gould, and Robert L Yauch. Prc2-mediated repression of smarca2 predicts ezh2 inhibitor activity in swi/snf mutant tumors. Proceedings of the National Academy of Sciences, 114(46):12249–12254, 2017.

Aimable Robert Jonckheere. A distribution-free k-sample test against ordered alternatives. Biometrika, 41 (1/2):133–145, 1954.

Daesik Kim, Sangsu Bae, Jeongbin Park, Eunji Kim, Seokjoong Kim, Hye Ryeong Yu, Jinha Hwang, Jong-Il Kim, and Jin-Soo Kim. Digenome-seq: genome-wide profiling of crispr-cas9 off-target effects in human cells. Nature methods, 12(3):237, 2015.

Daesik Kim, Sojung Kim, Sunghyun Kim, Jeongbin Park, and Jin-Soo Kim. Genome-wide target specificities of crispr-cas9 nucleases revealed by multiplex digenome-seq. Genome research, 26(3):406–415, 2016.

Eiru Kim, Merve Dede, Walter F Lenoir, Gang Wang, Sanjana Srinivasan, Medina Colic, and Traver Hart. Hierarchical organization of the human cell from a cancer coessentiality network. bioRxiv, page 328880, 2018.

Ben Langmead, Cole Trapnell, Mihai Pop, and Steven L Salzberg. Ultrafast and memory-efficient alignment of short dna sequences to the human genome. Genome biology, 10(3):R25, 2009.

Yanni Lin, Thomas J Cradick, Matthew T Brown, Harshavardhan Deshmukh, Piyush Ranjan, Neha Sarode, Brian M Wile, Paula M Vertino, Frank J Stewart, and Gang Bao. Crispr/cas9 systems have off-target activity with insertions or deletions between target dna and guide rna sequences. Nucleic acids research, 42(11):7473–7485, 2014.

Yunhua Liu, Xinna Zhang, Cecil Han, Guohui Wan, Xingxu Huang, Cristina Ivan, Dahai Jiang, Cristian Rodriguez-Aguayo, Gabriel Lopez-Berestein, Pulivarthi H Rao, et al. Tp53 loss creates therapeutic vulnerability in colorectal cancer. Nature, 520(7549):697, 2015.

E Robert McDonald III, Antoine De Weck, Michael R Schlabach, Eric Billy, Konstantinos J Mavrakis, Gregory R Hoffman, Dhiren Belur, Deborah Castelletti, Elizabeth Frias, Kalyani Gampa, et al. Project drive: a compendium of cancer dependencies and synthetic lethal relationships uncovered by largescale, deep rnai screening. Cell, 170(3):577–592, 2017.

Robin M Meyers, Jordan G Bryan, James M McFarland, Barbara A Weir, Ann E Sizemore, Han Xu, Neekesh V Dharia, Phillip G Montgomery, Glenn S Cowley, Sasha Pantel, et al. Computational correction of copy number effect improves specificity of crispr-cas9 essentiality screens in cancer cells. Nature Genetics, 2017.

Huaiyu Mi, Xiaosong Huang, Anushya Muruganujan, Haiming Tang, Caitlin Mills, Diane Kang, and Paul D Thomas. Panther version 11: expanded annotation data from gene ontology and reactome pathways, and data analysis tool enhancements. Nucleic acids research, 45(D1):D183–D189, 2016.

David W Morgens, Michael Wainberg, Evan A Boyle, Oana Ursu, Carlos L Araya, C Kimberly Tsui, Michael S Haney, Gaelen T Hess, Kyuho Han, Edwin E Jeng, et al. Genome-scale measurement of off-target activity using cas9 toxicity in high-throughput screens. Nature communications, 8:15178, 2017.

Florian L Muller, Simona Colla, Elisa Aquilanti, Veronica E Manzo, Giannicola Genovese, Jaclyn Lee, Daniel Eisenson, Rujuta Narurkar, Pingna Deng, Luigi Nezi, et al. Passenger deletions generate therapeutic vulnerabilities in cancer. Nature, 488(7411):337, 2012.

Diana M Munoz, Pamela J Cassiani, Li Li, Eric Billy, Joshua M Korn, Michael D Jones, Javad Golji, David A Ruddy, Kristine Yu, Gregory McAllister, et al. Crispr screens provide a comprehensive assessment of cancer vulnerabilities but generate false-positive hits for highly amplified genomic regions. Cancer discovery, 6 (8):900–913, 2016.

Deepak Nijhawan, Travis I Zack, Yin Ren, Matthew R Strickland, Rebecca Lamothe, Steven E Schumacher, Aviad Tsherniak, Henrike C Besche, Joseph Rosenbluh, Shyemaa Shehata, et al. Cancer vulnerabilities unveiled by genomic loss. Cell, 150(4):842–854, 2012.

Henriette O’Geen, Isabelle M Henry, Mital S Bhakta, Joshua F Meckler, and David J Segal. A genome-wide analysis of cas9 binding specificity using chip-seq and targeted sequence capture. Nucleic acids research, 43(6):3389–3404, 2015.

H Pagés. Bsgenome: Infrastructure for biostrings-based genome data packages and support for efficient snp representation. R package, 2016.

Inju Park, Cecil Han, Sora Jin, Boyeon Lee, Heejin Choi, Jun Tae Kwon, Dongwook Kim, Jihye Kim, Ekaterina Lifirsu, Woo Jin Park, et al. Myosin regulatory light chains are required to maintain the stability of myosin ii and cellular integrity. Biochemical Journal, 434(1):171–180, 2011.

José Manuel Pérez-Pérez, Héctor Candela, and José Luis Micol. Understanding synergy in genetic interactions. Trends in genetics, 25(8):368–376, 2009.

Ophir Shalem, Neville E Sanjana, and Feng Zhang. High-throughput functional genomics using crispr–cas9. Nature Reviews Genetics, 16(5):299, 2015.

Teunis J Terpstra. The asymptotic normality and consistency of kendall’s test against trend, when ties are present in one ranking. Indagations Math., 14:327–333, 1952.

Shengdar Q Tsai and J Keith Joung. Defining and improving the genome-wide specificities of crispr–cas9 nucleases. Nature Reviews Genetics, 17(5):300, 2016.

Shengdar Q Tsai, Zongli Zheng, Nhu T Nguyen, Matthew Liebers, Ved V Topkar, Vishal Thapar, Nicolas Wyvekens, Cyd Khayter, A John Iafrate, Long P Le, et al. Guide-seq enables genome-wide profiling of off-target cleavage by crispr-cas nucleases. Nature biotechnology, 33(2):187, 2015.

Aviad Tsherniak, Francisca Vazquez, Phil G Montgomery, Barbara A Weir, Gregory Kryukov, Glenn S Cowley, Stanley Gill, William F Harrington, Sasha Pantel, John M Krill-Burger, et al. Defining a cancer dependency map. Cell, 170(3):564–576, 2017.

Tim Wang, Kıvan¸c Birsoy, Nicholas W Hughes, Kevin M Krupczak, Yorick Post, Jenny J Wei, Eric S Lander, and David M Sabatini. Identification and characterization of essential genes in the human genome. Science, 350(6264):1096–1101, 2015.

Han Xu, Tengfei Xiao, Chen-Hao Chen, Wei Li, Clifford A Meyer, Qiu Wu, Di Wu, Le Cong, Feng Zhang, Jun S Liu, et al. Sequence determinants of improved crispr sgrna design. Genome research, 25(8):1147–1157, 2015.

